# Comparison of genetic variation between rare and common congeners of *Dipodomys* with estimates of contemporary and historical effective population size

**DOI:** 10.1101/2021.08.04.455110

**Authors:** Michaela K. Halsey, John D. Stuhler, Natalia Bayona-Vasquez, Roy N. Platt, Jim R. Goetze, Robert E. Martin, Kenneth G. Matocha, Robert D. Bradley, Richard D. Stevens, David A. Ray

**Affiliations:** Department of Biological Sciences, Texas Tech University, Lubbock, Texas, United States of America; Department of Natural Resources Management, Texas Tech University, Texas, United States of America; Department of Environmental Health Science, University of Georgia, Athens, Georgia, United States of America; Institute of Bioinformatics, University of Georgia, Athens, Georgia, United States of America; Texas Biomedical Research Institute, San Antonio, Texas, United States of America; Natural Sciences Department, Laredo Community College, Laredo, Texas, United States of America; Department of Biology, McMurry University, Abilene, Texas, United States of America; Department of Biology, South Arkansas Community College, El Dorado, Arkansas, United States of America; Natural Science Research Laboratory, Museum of Texas Tech, Lubbock, Texas, United States of America

**Keywords:** 3RAD, *Dipodomys elator*, *Dipodomys ordii*, genetic structure, metapopulation, single nucleotide polymorphisms

## Abstract

Organisms with low effective population sizes are at greater risk of extinction because of reduced genetic diversity. *Dipodomys elator* is a kangaroo rat that is classified as threatened in Texas and field surveys from the past 50 years indicate that the distribution of this species has decreased. This suggests geographic range reductions that could have caused population fluctuations, potentially impacting effective population size. Conversely, the more common and widespread *D. ordii* is thought to exhibit relative geographic and demographic stability. Genetic variation between *D. elator* and *D. ordii* samples was assessed using 3RAD, a modified restriction site associated sequencing approach. It was hypothesized that *D. elator* would show lower levels of nucleotide diversity, observed heterozygosity, and effective population size when compared to *D. ordii*. Also of interest was identifying population structure within contemporary samples of *D. elator* and detecting genetic variation between temporal samples that could indicate demographic dynamics. Up to 61,000 single nucleotide polymorphisms were analyzed. It was determined that genetic variability and effective population size in contemporary *D. elator* populations were lower than that of *D. ordii*, that there is only slight, if any, structure within contemporary *D. elator* populations, and there is little genetic differentiation between spatial or temporal historical samples suggesting little change in nuclear genetic diversity over 30 years. Results suggest that genetic diversity of *D. elator* has remained stable despite claims of reduced population size and/or abundance, which may indicate a metapopulation-like system, whose fluctuations might counteract any immediate decrease in fitness.

## Introduction

Measuring genetic variation in rare, threatened, endemic, or endangered species has important implications for management and is integral to conservation efforts [1]. Population genetic summary statistics can be used to delimit management units based on significantly different allele frequencies [2], identify population structure [3], or assess connectivity of demographically disparate subpopulations [4]. One such critical measure for small populations is effective population size, N_e_ [5]. Effective population size can be influenced by fluctuations in census size, mating strategy, biased sex ratios, migration, demographic history, spatial dispersion, and population structure [6-9] and typically is far less than the census size. However, any one N_e_ value is hard to interpret because it lacks a context. One such context to help understand the potential impacts of fluctuations of N_e_ is comparison between more restricted, possibly threatened species and a widespread congener, which are presumed to harbor more genetic variation.

The Texas kangaroo rat (*Dipodomys elator*) is a heteromyid rodent that has a limited distribution in north-central Texas [10-14]. Though previously found in two counties in Oklahoma, it appears to have been extirpated from that state [15]. Moreover, *D. elator* has a small geographic range and low dispersal capability [16-17], which increases isolation from nearby subpopulations.

The distribution of the Texas kangaroo rat appears dynamic [18]. For instance, though the species was described from a specimen in Clay County [19], it has not been observed there in more recent surveys. Additionally, resampling of sites where it has been previously documented have failed to detect the species, and new localities of presence have been identified in more recent surveys [20]. Furthermore, previous studies of *D. elator* population genetics [21-22] have been relevant to assess genetic diversity and structure within the species and have established a valuable reference point for *D. elator*. However, these studies relied on few molecular markers (i.e. enzymes, mitochondrial DNA, and microsatellites). Additionally, it is useful to compare contemporary samples to historical collections to identify potential changes in overall diversity [23-24].

Ord’s kangaroo rat, *D. ordii* is a medium sized rodent that occurs from Canada into Mexico [25] Given its large geographic range and preferred commonly available habitat choice (i.e. sandy soils), *D. ordii* is not listed on any state or U.S. federal critically threatened and endangered lists. The population in Canada, however, is listed as endangered [26]. To our knowledge, there has not been a range-wide genetic analysis of *D. ordii*, and the last regional genetic study on *D. ordii* isoenzymes was published by [27].

Here, we compare population genetic parameters of *D. elator* with *D. ordii*, further compare *D. elator* samples from two time periods (pre- and post-2000), and for contemporary samples, investigate differences in genetic diversity across the distribution. We make several predictions: 1) *D. elator* will exhibit a lower effective population size than *D. ordii*, and concomitantly, lower nucleotide diversity, lower observed heterozygosity, and higher inbreeding coefficients; 2) there will be greater genetic diversity among contemporary samples of *D. elator* than in historical samples, as contemporary samples were taken from across the distribution, compared to historical samples collected from three counties in the middle of its distribution; and 3) historical N_e_ from a coalescent approach for *D. ordii* and *D. elator* will demonstrate that *D. elator* exhibits a lower N_e_ at present than *D. ordii*.

## Methods

### Sample collection

Kangaroo rats were captured using Sherman live traps (23×9×8 cm; H.B. Sherman Traps, Inc. Tallahassee, Florida) during surveys within the historical range of *D. elator* (Fig 1) from 2015 to 2017. When a *D. elator* individual was captured, it was either 1) taken as a voucher specimen for deposition at the Natural Science Research Laboratory (NSRL) at the Museum of Texas Tech University or, 2) had between two to four whiskers extracted from either side of the rostrum [28]. In the latter case, thicker whiskers (i.e., macrovibrissae) were selected with the follicle intact. Whiskers were stored in a sterile vial with 1% sodium dodecyl sulfate (SDS) lysis buffer [29]. A buccal swab was also collected from one *D. elator* individual as described in detail in [28].

**Fig 1.**
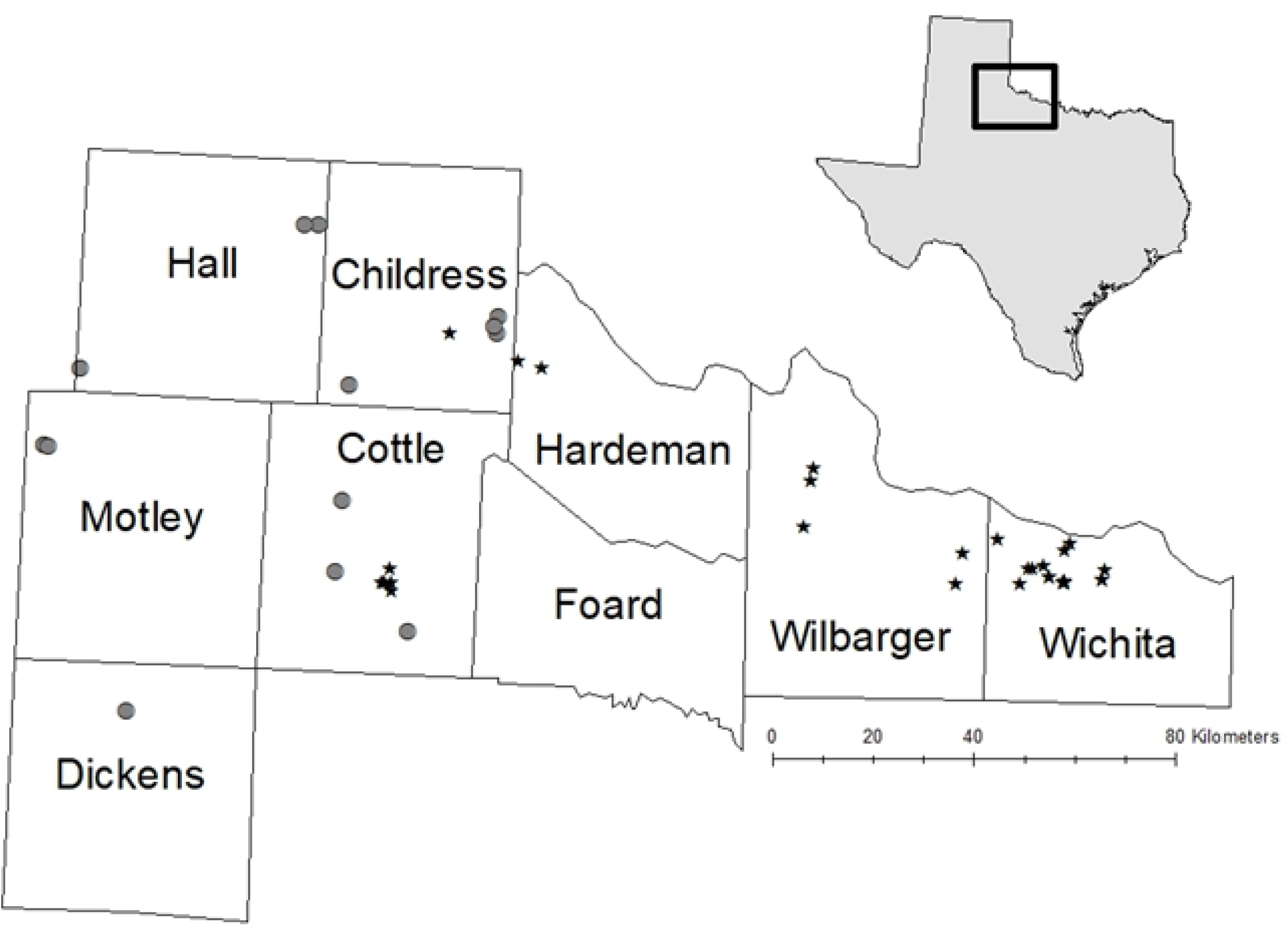
Map of contemporary kangaroo rat samples used in this study. Filled stars indicate *Dipodomys elator* samples whereas gray circles represent *D. ordii* samples used in the study. Note the sampling hole located in Foard County, most of Hardman County, and in south Wilbarger County. Trapping restrictions and topography prevented collections in those regions.

Other methods of collecting DNA from rats included tail salvages and from toe clips from museum specimens (Table S1). *D. elator* tail lengths average about 196 mm [30] and at times the end of the tail (i.e., the plume) was severed by the door of an activated Sherman trap. These salvaged tail plumes were placed in sterile vials of 1% sodium dodecyl sulfate (SDS) lysis buffer [29]. Also sampled were toe clips that had been collected from rats from 1986 to 1995 as part of a genetic survey of the species by REM and KGM.

In total, 70 *D. elator* samples were analyzed from five tissue types (i.e, liver, whisker, tail, buccal swab, and toe clips) and two time periods (prior to 2000 and contemporary surveys from 2015 to 2017; Table S1). Additionally, 26 *D. ordii* liver samples were collected in five counties from 2015 to 2017. Contemporary sampling followed guidelines established by the American Society of Mammalogists [31]. Animal handling protocols were approved by the Institutional Animal Care and Use Committee at Texas Tech University (#T14083).

Throughout the manuscript the *D. elator* samples will be referred to using the following descriptors: ‘historical’, collected prior to 2000; ‘contemporary’, collected after 2000; ‘west’, collected from Cottle, Childress, or Hardeman counties; and ‘east’ collected from Baylor, Wilbarger and Wichita counties (Fig 1).

### DNA extraction

DNA was extracted using the Qiagen DNeasy Blood and Tissue spin column protocol (Qiagen; Venlo, Netherlands). For liver, toe clips, and tail salvages, the manufacturer’s recommendations were followed. For whisker and buccal swab samples, the protocol found in [28] was implemented. In all cases, DNA concentration was fluorometrically quantified using the Qubit 3.0, high sensitivity assay (Invitrogen, Life Technologies, Carlsbad, CA).

### 3RAD library prep, sequencing, and data husbandry

RADseq libraries were prepared following the 3RAD protocol found in [32] Details of library prep conditions used in this study are provided in Supplemental File S1. In short, restriction enzyme combinations were tested in a subset of samples from both species, and according to digestion patterns and pilot sequence data, the best combination (i.e. MspI, EcoRI, and ClaI), was further used for all samples. Samples were normalized, digested, enzyme-specific adapters were ligated, and ligation products were purified. To generate full-length library constructs, ligated products were amplified using iTru5 and iTru7 primers [33]. For most samples, a molecular ID tag (iTru 5 8N) was incorporated in the first cycle of PCR, to detect PCR duplicates [32, 34]. PCR products were purified, pooled, and size-selected at a range of 550 bp +/- 15%. Size-selected fragments were purified and sequenced using an Illumina HiSeq to generate paired end data at Oklahoma Medical Research Foundation Genomics Core or Novogene Inc.

Stacks v1.48 and v2.01 [35] was employed to demultiplex, analyze, and export data into other formats. After demultiplexing, poor reads were filtered using the AfterQC ‘after.py’ pipeline [36]. Poor reads were defined as exhibiting a low quality score (PHRED score < 15), bad overlaps (i.e., mismatched reads), too many ambiguous nucleotides (greater than 40% of the read), short read lengths (< 35 base pairs), or homopolymer regions. If a read failed one of these steps, it was removed from downstream analyses. Reads were aligned within Stacks to the *D. ordii* genome assembly (accession ID GCA_000151885.1) using the Burrows-Wheeler aligner [37].

Data were grouped into putative loci, and polymorphisms were identified with the ‘gstacks’ module in Stacks. Common population genetic statistics such as observed and expected heterozygosity, nucleotide diversity, and inbreeding coefficients were calculated using the ‘populations’ module. This step was repeated four times to determine a balance between data used and processing speed. Though the “gold standard” is to include loci where 75 to 80% of the individuals in a population have that locus, known as the -r value [35, 38], this has been shown to bias population genetic measures, especially in cases where data are not plentiful. This influences biological implications [39-42]. The ‘populations’ module using this 75% rule (-r 0.75), two liberal filters (-r of 0.25 and 0.5) and a more conservative filter (-r 0.95) was run. For most downstream analyses, -r 0.75 was used as the main dataset and for comparison across -r values for any differences. Finally, loci and individuals that had greater than 20% missing data were removed.

### Population genetics

Observed and expected heterozygosity were calculated using the ‘summary’ function in the R package adegenet, version 2.1.1 [43]. F_IS_ and F_ST_ values were calculated using hierfstat [44]. Nei’s genetic distances [45] were determined and plotted using the ‘aboot’ function in the poppr R package [46].

### Estimation of effective population size

To determine effective population size using NeEstimator v2.1 [51], the Genepop file [52] generated by Stacks was used on our contemporary dataset. NeEstimator calculates Ne using three methods: linkage disequilibrium, molecular co-ancestry, and a temporal method. The first two methods were used to determine contemporary effective population sizes per species.

For historical Ne of *D. elator*, the Extended Bayesian Skyline Plot (EBSP) coalescent test as implemented in BEAST 2.0 [53] was used. Once loci containing multiple single nucleotide polymorphisms were determined, the protocol outlined in [54] was followed, using a strict molecular clock set to 1.0 and a generation time of 3 years [55]. Plots were constructed with 24 individuals and 47 loci.

This same process was followed for *D. ordii*; however, 15 individuals and 49 loci were used. Only one individual from Dickens County (Fig 1) was included to avoid misinterpretations due to possible inbreeding since many individuals collected from that county were collected from a single location.

### Population structure

To infer population structure for each species, the STRUCTURE algorithm was used [47]. All singletons and private doubletons were removed, which have been shown to mask weak population structure [48-49]. Only one randomly selected SNP from each locus was used to minimize possible effects of linked data. For all runs, 50,000 burn-in iterations were executed and 200,000 Markov Chain Monte Carlo (MCMC) repetitions with 3 replicates at each K, which ranged from 1 to 5. The program DISTRUCT v1.1 was used to visualize the final output of structure analyses [50].

### Principal Components Analysis

To visualize genetic structure of the population without assigning individuals to clusters *a priori*, a Principal Components Analysis was conducted using the function dudi.pca in the R package ‘adegenet’ version 2.1.1 [43] on historical and contemporary samples. Only the first two axes were retained for all datasets based on the scree plots generated by glPca.

## Results

In all, 96 kangaroo rats were sequenced and analyzed from two species in eight counties in north-central Texas. 3RAD analysis for 70 individual *D. elator* samples produced over 34 million reads. Before filtering within the ‘populations’ module, there were 330,326 loci suitable for analysis. Approximately 1.5% of reads per sample were removed from the dataset following AfterQC filtering. Similar analysis for the 26 *D. ordii* samples produced over 10 million reads. Prior to ‘populations’ filtering, approximately 382,514 loci were eligible for further analysis. Fewer than 2% of reads per sample were removed from the dataset. For *D. elator* samples, after removing loci and individuals that had greater than 20% missing data, 3.935 samples remained from 55 individuals.

### Summary population genetics

There were as few as 7 single nucleotide polymorphisms (SNPs) to as many as 61,000 SNPs to analyze (Table S2). The general trend was that there were fewer SNPs analyzed as -r value increased and extreme values of -r (i.e. 0.25 and 0.95) yielded stronger deviations between observed and expected heterozygosity across all analyses (Table S2).

When compared to each other, contemporary *D. elator* samples showed lower levels of observed heterozygosity (0.042-0.043) than *D. ordii* (0.128) which suggests lower genetic diversity. F_IS_ was positive in all *D. elator* groups (0.015-0.031) except for in historical samples (−0.007). F_ST_ values show moderately low genetic differentiation (0.035-0.041) across *D. elator* comparisons (Table 1).

**Table 1.**
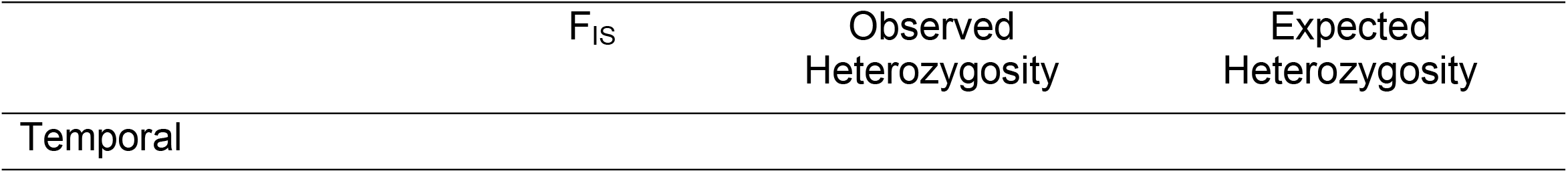
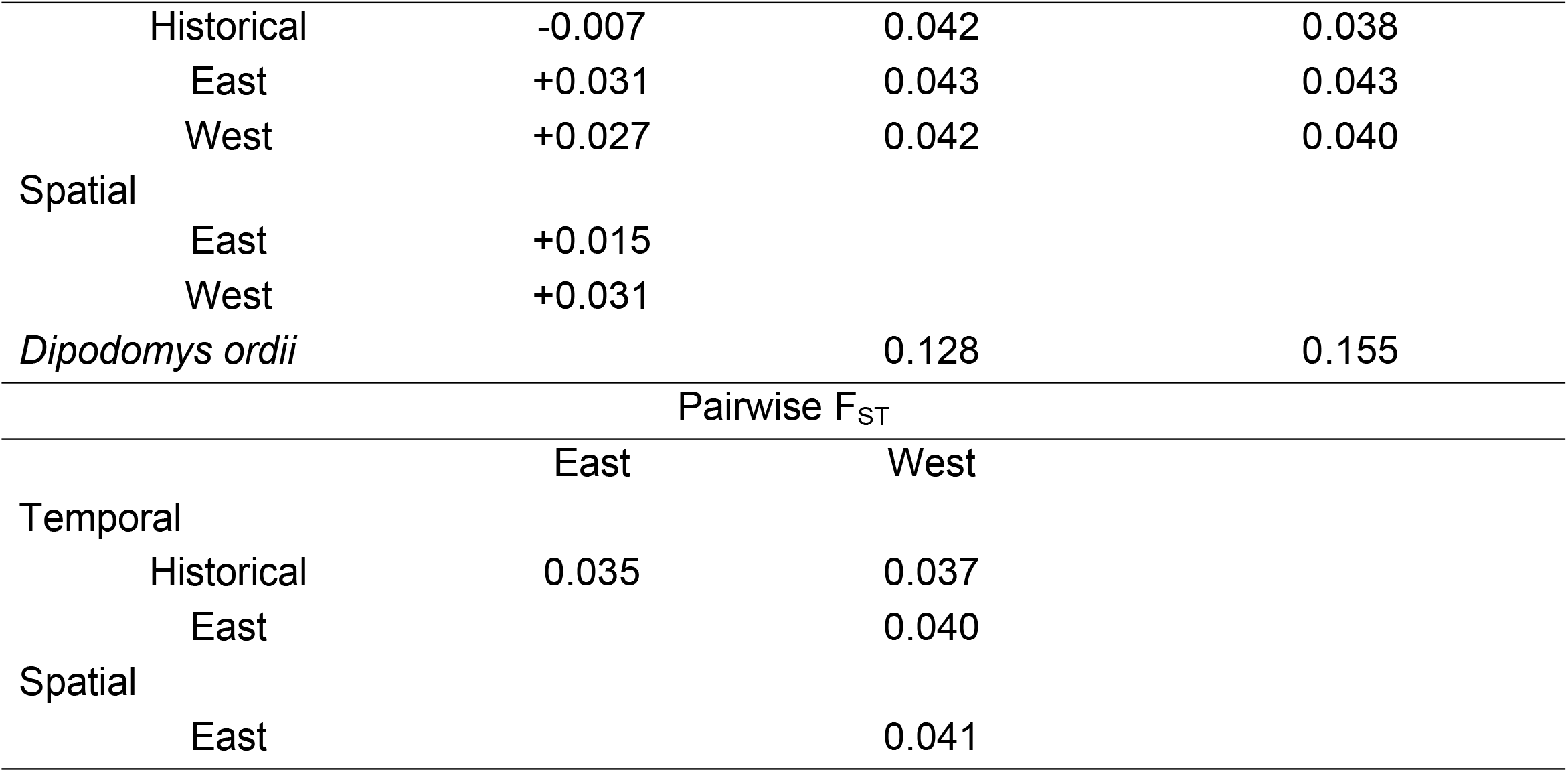
Genetic diversity summary statistics for *D. elator* and observed heterozygosity and expected heterozygosity values for *D. ordii*. Only one population is assumed for *D. ordii*, so there are no values for F_ST_ or F_IS_.

| | $F_{IS}$ | Observed<br>Heterozygosity | Expected<br>Heterozygosity |
| --- | --- | --- | --- |
| Temporal |  |  |  |

|  |  |  |  |  |
| --- | --- | --- | --- | --- |
|  | Historical | -0.007 | 0.042 | 0.038 |
|  | East | +0.031 | 0.043 | 0.043 |
|  | West | +0.027 | 0.042 | 0.040 |
| Spatial | East | +0.015 |  |  |
|  | West | +0.031 |  |  |
| *Dipodomys ordii* |  |  | 0.128 | 0.155 |
| Pairwise $F_{ST}$ | | | | |
|  |  | East | West |  |
| Temporal |  |  |  |  |
|  | Historical | 0.035 | 0.037 |  |
|  | East |  | 0.040 |  |
| Spatial |  |  |  |  |
|  | East |  | 0.041 |  |

### Current and historical effective population size

Only the linkage disequilibrium method in NeEstimator v.2.1 produced a value other than ‘Infinite’ for effective population size for *D. elator*. Estimated N_e_ of the east group 171.3, with a 95% confidence interval of 158-186.9 using the lowest allele frequency. For the west group, both the linkage disequilibrium and molecular co-ancestry methods returned ‘Infinite’ for N_e_ at all allele frequencies. No method with NeEstimator was able to provide an estimate of population size for *D. ordii*, other than ‘Infinite.’

The Extended Bayesian Skyline plots generated for the *D. elator* dataset showed a decline in effective population size over the last 10,000 years, to an approximate current N_e_ of 500. For *D. ordii*, N_e_ has increased in the last 5,000 years and is estimated to stand at about 10,000 individuals (Fig 2).

**Fig 2.**
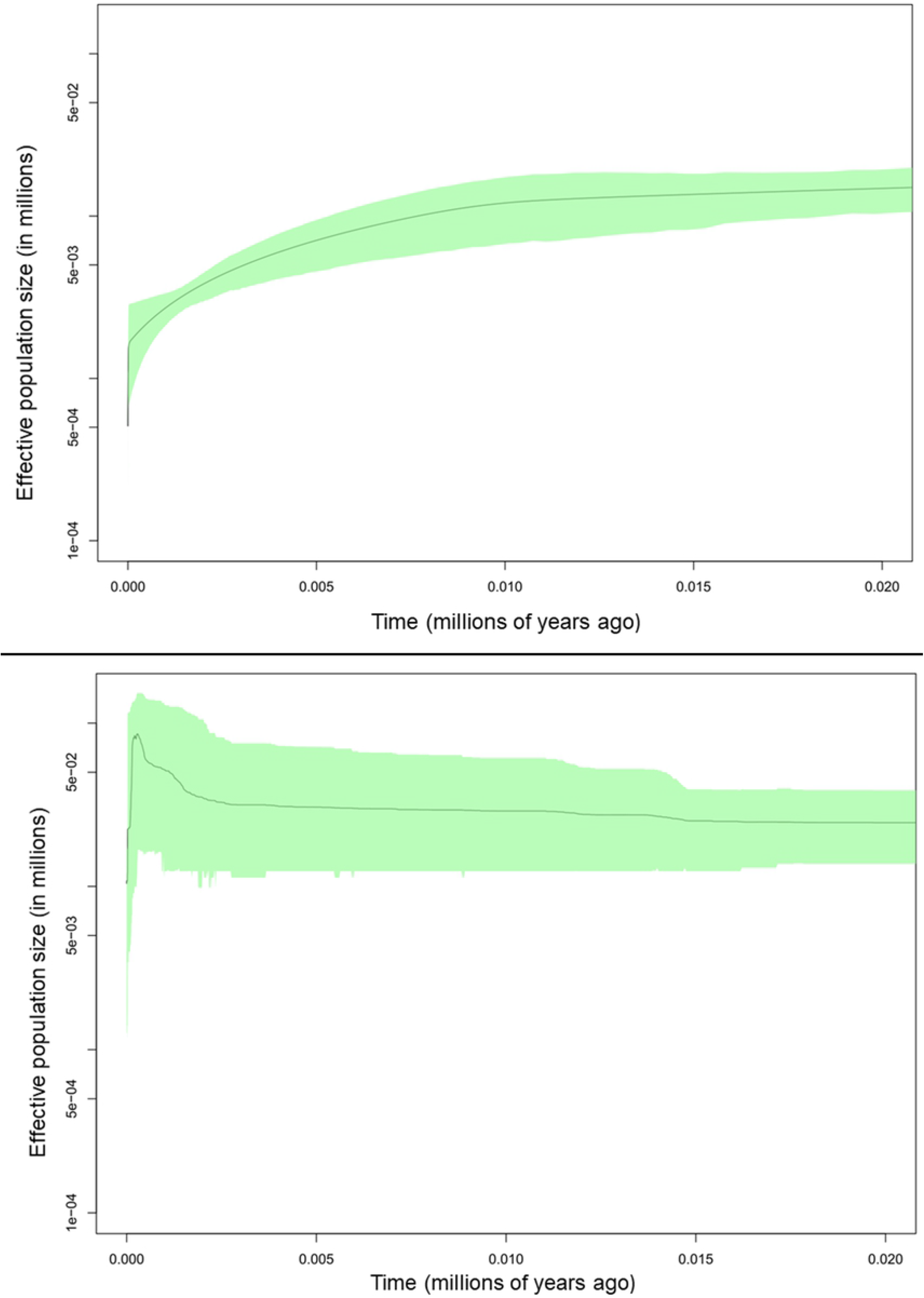
Extended Bayesian Skyline Plot for *Dipodomys elator* (top) from 34 individuals and 47 SNPs and for *D. ordii* (bottom) from 15 individuals and 49 SNPs. X-axis is millions of years ago. Y-axis is effective population size (Ne) in millions.

### Population substructure

Based on log-likelihood scores (Table S3) and their respective variances from STRUCTURE, the “best” value for k for *D. elator* was 3. When visualizing the PCA bi-polot, PC1 accounts for almost 98% of the variation found in the dataset and shows geographic separation along PC2, which only accounts for 0.1% of the variation (Fig 3). Using Nei’s genetic distance, most contemporary samples from the west cluster together and are nested within historical samples (Fig 4). The historical PCA for *D. elator* samples excluding those from the 1960s (Fig S1) largely confirms that all individuals were taken from the same region (Hardeman County).

**Fig 3.**
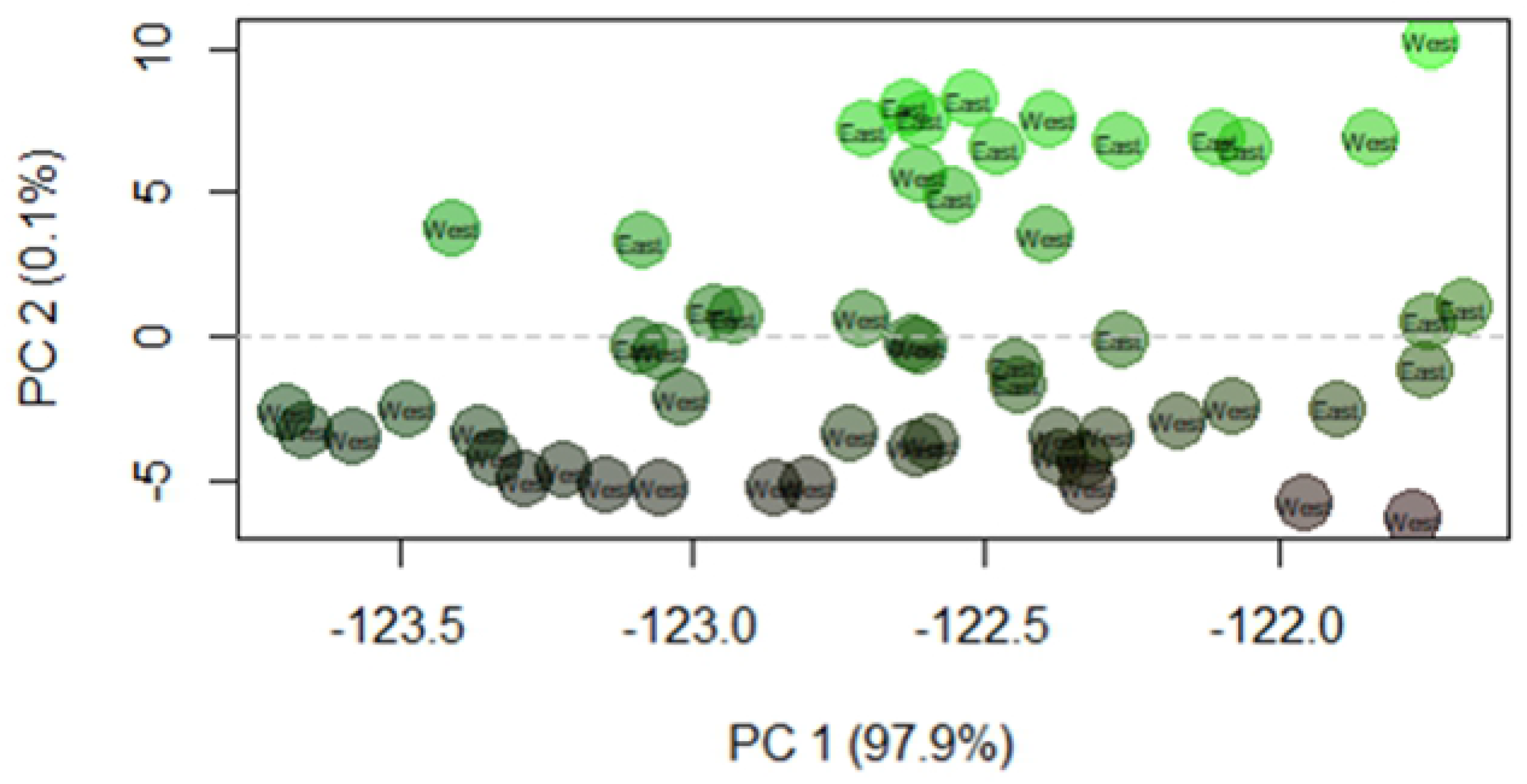
Principal components analysis on the genotypes for 55 samples (historical and contemporary) using the dudi.pca function in R package ‘adegenet’. While there are no clear clusters emerging on PC1, geographic location seems to correspond with PC2.

**Fig 4.**
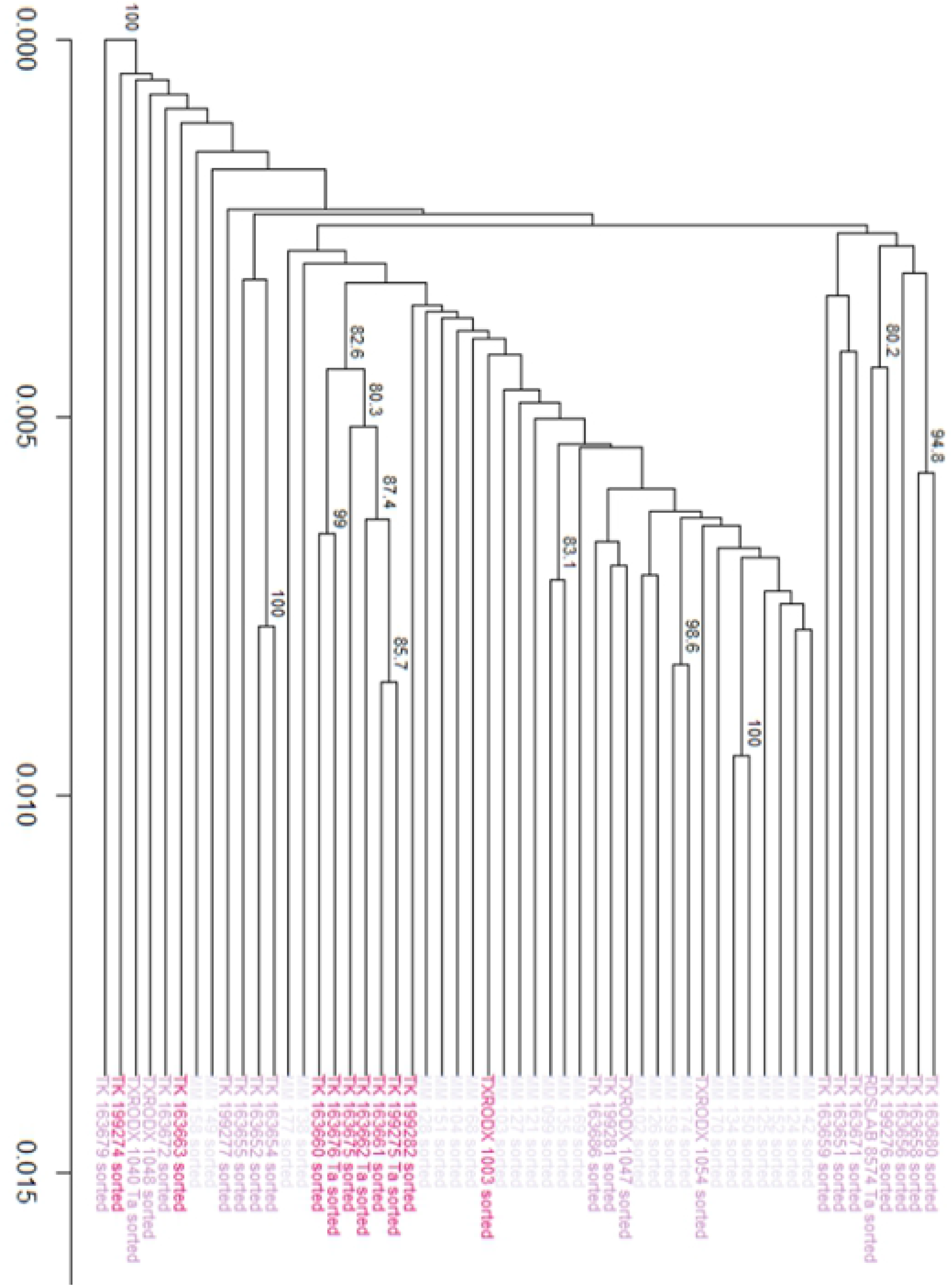
Nei’s genetic distance dendrogram for 55 samples (historical and contemporary). The patchy arrangement of individuals suggests gene flow between the hypothesized east and west populations.

## Discussion

This study evaluated changes in genetic diversity across time and space by comparing a rare species with a hypothesized amorphous and restricted distribution to a more common congener with a larger, more defined range. This is only the third population genetic study on *Dipodomys elator* in over 30 years and it is the first to make use of genomic techniques, screening from tens to thousands of markers, making the study valuable for current and future conservation efforts. In [21], allozyme markers were used to conclude that there was moderate genetic differentiation among three *D. elator* localities (Hardeman, Wilbarger, and Wichita counties). This is seemingly incongruent with our results in which we observed little genetic differentiation (F_ST_ = 0.041), but the difference could simply be the result of the markers used (SNPs versus allozymes).

More recently, [22] observed low mitochondrial DNA variation but high microsatellite diversity within the species. They concluded that genetic drift and not gene flow has had a greater impact on configuring *D. elator* genetic diversity. This result is possible because mitochondrial DNA has a lower effective population size than neutral nuclear markers such as RAD loci [56]. Genetic drift could play a role in structuring mitochondrial DNA diversity, but more time would be needed to detect reduction of diversity in the nuclear genome using older markers such as microsatellites. An insufficient number of polymorphic microsatellite loci limits genetic resolution between individuals with supposed low population-level diversity. Our results suggest that RAD loci, that have a slower rate of mutation than microsatellites, are superior when investigating populations with weak population structure [57]. Finally, genomic data generated from this study can be used for future genomic investigations, such as those examining family structure [58].

Together, these three studies, using allozyme, mtDNA, microsatellite and RAD-Seq markers, offer numerous mean geographic estimates of F_ST_ within this species. In [21], the mean F_ST_ was found to be 0.102, [22] estimated F_ST_ of 0.096 from their late 1960s samples, and our study, at the greatest resolution of all previous studies, reveals a mean F_ST_ value of 0.041. Our lower mean value includes individuals sampled from localities (Cottle and Childress counties) not present in the previous two studies. These results suggest modest population differentiation corresponding with geography.

From an overall genetic diversity perspective, our data suggest that there has not been a substantial loss in genetic diversity over the last 30 years, despite what seems to be a decrease in the distribution (and possibly abundance) of *Dipodomys elator*, similar to what [59] found in *D. ingens*, the giant kangaroo rat. In other words, despite a decline in distribution and census size, the genetic diversity of the species is sufficiently high to offset any short-term effects of reduced fitness. This is supported by our estimate of N_e_ of between 170 to 500, which exceeds the recommended value to curtail inbreeding depression, as outlined by the 100/1000 rule [60]. Within this range, there exist enough individuals to mitigate immediate reduction in fitness but is not sustainable in the longer term (over thousands of years). In [22], it was also found that the N_e_ of this species was between 65 and 490 individuals.

Results from our contemporary samples confirm that subpopulation differentiation is not substantial (F_ST_ < 0.05). The STRUCTURE algorithm determined the best value of k to be 3. However, examining the plots suggests that samples represent a single interbreeding population. More clusters (i.e. subpopulations), while possible, are not biologically practical. This may just be an artefact of our sampling scheme (for example, k=5, one for each county). Second, newly colonized subpopulations on the fringes of ranges can exhibit lower levels of genetic diversity than expected [61]. For our contemporary samples, this is not the case; the low mean value of F_ST_ (< 0.05) does not seem to support cluster sizes of k=4 and k=5. Based on climate, vegetation, edaphic, and land use characteristics across the study area [20], our *a priori* assumption was that there are two subpopulations (east and west). However, STRUCTURE, the PCA, and Nei’s genetic distances do not clearly support two distinct subpopulations, suggesting there is a fair amount of gene flow in the region.

Our *a priori* subpopulations display low levels of inbreeding and very little genetic differentiation, suggesting one large interbreeding population (though not necessarily panmictic). Our samples were collected on opposite sides of a cline, separated by a region of inaccessible private land, so it was difficult to determine if the slight differentiation is due to that distance or if there is true population substructure and isolation from other habitat patches [62]. We included additional historic samples from specimens collected in the 1960s from areas within this “sampling hole” to answer this question. We anticipated that if the contemporary east and west subpopulations were indeed distinct, then genetic differentiation would be greater between them than to the samples from the sampling hole. In other words, a STRUCTURE plot would show the sampling hole samples as intermediate between the two. Alternatively, if the contemporary east and west subpopulations were considered one population then we would expect greater genetic differentiation between them and our “sampling hole” samples. Our results support the second prediction (Fig S2). However, the periods separating the datasets (anywhere between 20 and 50 years) and the relatively short generation times of kangaroo rats (about 3 years; Pacifici et al. 2012) would lead to high genetic turnover, so these results must be interpreted with caution. If there is substantial genetic turnover, this too could indicate small current effective population size, which supports our estimate of approximately 171.

As expected, our *D. ordii* samples exhibited higher genetic diversity estimates in nearly all categories despite our samples being collected from only five counties in north-central Texas. This emphasizes the substantial genetic diversity and evolutionary potential displayed by the common *D. ordii*, compared with a much rarer congener. However, we were surprised to find that *D. ordii* had a greater inbreeding coefficient than *D. elator* across some analyses. This pattern can be attributed to sampling bias, given that we sampled from a small portion of the *D. ordii* range, and half of the *D. ordii* samples were collected from a single ranch in Dickens County, Texas, where most individuals collected may present a certain degree of relatedness by proximity. Comparing between individuals from this ranch and a similarly situated subset of *D. elator* individuals, expected heterozygosity, π, and inbreeding coefficients were largely similar (Table S4). This suggests that potentially related individuals of *D. elator* do not show reduced genetic diversity than similarly related *D. ordii* individuals.

We were unable to generate a value of N_e_ for the current sample of *D. ordii*, likely a result of samples displaying high degrees of relatedness, so we used the value calculated from EBSP, which was approximately 10,000 individuals. In contrast to *D. elator* N_e_, which declined over time, the plot for *D. ordii* increased, perhaps an indication of colonization of new habitat (northward) as glaciers receded after the Last Glacial Maximum 20,000 years ago [63-64].

Coupled with low N_e_ estimates, and population surveys that recover or fail to locate *D. elator* in different localities, one possibility is that this population exhibits characteristics of a metapopulation [65-66]. Metapopulation theory has been discussed in the context of mammalian conservation biology because it accommodates populations in fragmented habitats [67], but empirical studies to develop metapopulation theory for threatened and endangered mammals are few (see [68-69]). One reason for the difficulty to meet the original metapopulation criteria of [70] is the stringency of the original criteria. In [71], the authors relaxed two criteria, adding that subpopulations, not the colonized habitat patch, are the discrete entity, and that these discrete subpopulations differ in their demography, implying asynchronicity. Based on field surveys, analysis of field notes, museum specimens, and species distribution models [20] there is evidence that the *D. elator* population may benefit from management consideration stemming from metapopulation theory.

However, because this connection to metapopulation theory is still tenuous, the overall population should still be monitored [72]. Perhaps a long-term demographic study is warranted. Managing the metapopulation must be concerned with maintaining dispersal and gene flow and other population dynamics among subpopulations. Should managers elect for extreme measures to manage *D. elator* populations, such as translocations or reintroductions, knowledge that the population is a metapopulation is critical. Lastly, it is important to note that the metapopulation in a conservation context has several assumptions. One assumption is the “equilibrium” between colonization and extinction across long time scales (i.e. if one patch goes extinct, another is colonized). This seems unlikely in many natural populations [73], including that of *D. elator*, but this type of assumption can be used to appropriately model changes in demography and genetics of *D. elator*.

There is no lack of research on habitat associations, mainly those evaluating soil and vegetation changes, as they influence *D. elator*. These studies have greatly improved our understanding of this elusive rodent [16-18, 74-75], but we still do not have answers to many basic biological questions. We do know, however, that the population of *D. elator* seems to track favorable habitat, albeit in a more restricted range than previously recorded [18]. Overall, the population of *D. elator* exhibits genetic variation lower than that of a species with a predictably greater effective population size. However, contemporary samples show no substantial decrease in genetic diversity from historical samples, suggesting that the *D. elator* population, though small and constantly shifting, has managed to maintain its genetic diversity.

This study demonstrates the effectiveness of using samples from gradations across the range, rather than at two extremes. Sampling from the extremes of a population range could lead researchers to inappropriate conclusions that could wrongly influence management decisions. Though the current effective population size of *D. elator* is estimated to be around 171 to 500 individuals, perhaps small population sizes are the status quo for this species. Increasing population size may be unsustainable for this species (greater competition, reduced resources, delayed or forgoing reproduction).

### Conservation Implications

Researchers interested in natural genetic variation and population structure of mammals should consider the possibility the population of their organisms of study could be exhibiting a metapopulation. This is especially important for species that are rarely seen or captured. Our findings suggest that the *D. elator* population could be a metapopulation that must be vigorously monitored so that managers can detect any great losses in genetic diversity and evolutionary potential. Furthermore, given the current advances in molecular techniques and analyses, it is no longer necessary to limit samples in the temporal dimension. Doing so, especially for species that remain understudied, will prove detrimental to any plan long-term plan for management. We advise continued use of reduced representation sequencing (ddRAD, 3RAD) but with inclusion of historic and geographically represented samples to fully encapsulate temporal and spatial genetic variability within a possibly imperiled species.

## CRediT authorship contribution statement

**Michaela Halsey:** Writing – Original Draft, Writing – Review and Editing, Formal Analysis, Investigation, Data Curation, Validation, Methodology, Visualization. **John Stuhler:** Methodology, Investigation, Writing – Review and Editing. **Natalia Bayona-Vasquez:** Methodology, Investigation, Data Curation, Writing – Original Draft, Writing – Review and Editing, Visualization. **Roy Platt II:** Conceptualization, Funding Acquisition, Methodology, Writing – Review and Editing. **Jim Goetze:** Conceptualization, Writing – Review and Editing. **Robert Martin:** Resources. **Kenneth Matocha;** Resources. **Robert Bradley:** Conceptualization, Methodology, Funding Acquisition, Writing -Review and Editing. **Richard Stevens:** Conceptualization, Methodology, Resources, Supervision, Project Administration, Funding Acquisition, Writing – Review and Editing. **David Ray:** Conceptualization, Methodology, Resources, Supervision, Project Administration, Funding Acquisition, Writing – Review and Editing

## Acknowledgements

Samples analyzed were loaned from the following collections: Museum of Texas Tech in Lubbock, Texas, Midwestern State University in Wichita Falls, Texas, and the Museum of Southwestern Biology in Albuquerque, New Mexico. Moreover, we would like to acknowledge Travis Glenn for suggesting 3RAD sequencing. This work would not be possible without cooperation from private landowners who allowed us to collect on their land. We sincerely appreciate the efforts of C. Garcia, C. Brothers, A. Kildow, J. Keats, S. de la Piedra, M. Krishnamoorthy, C. Rios-Blanco, G. Langlois, E. Stukenholtz, D. Arenas, L. Lindsay, T. Soniat, E. Roberts, E. Wright, I. Vasquez, J. Francis and many others for their help with fieldwork. Also, we would like to acknowledge the High-Performance Computing Center (HPCC) at Texas Tech University for providing substantial logistical support. We also like to thank H. Wilson and J. Grimshaw and all anonymous reviewers for constructive feedback used to improve earlier versions of this manuscript.

## Supporting Information

**Fig S1. Principal components analysis on the genotypes for historical samples from Hardeman County using the glPCA function in R package ‘adegenet’**.

**Fig S2. STRUCTURE plot of 70 D. elator samples across three time periods (see text for time breakdown)**. The “sampling hole” individuals (blue) are completely divergent from later samples; however, there is evidence of those loci persisting in the population. These results confirm a) no appreciable decrease in genetic diversity over 30 years and b) one interbreeding contemporary population).

**Table S1. Seventy *Dipodomys elator* samples and 26 *D. ordii* samples used in the genetic analysis including temporal (historical, contemporary) subpopulation, spatial (east or west) subpopulation, the specific county the individual was found, tissue type, and the museum where the voucher was received**. Museum codes are MSB (Museum of Southwestern Biology), MSU (Midwestern State University), and TTU (Texas Tech University).

**Table S2. Summary statistics calculated in Stacks for 26 *D. ordii* and 38 *D. elator* contemporary samples**. Private alleles are those alleles not shared with any other subpopulation.

**Table S3. Log-likelihood and delta K values used in the Evanno method for *D. elator* STRUCTURE analysis**.

**Table S4. General summary statistics calculated in Stacks for a comparison between 3 individuals from each species that were collected in proximity (i.e. same tract of land)**. Private alleles are those alleles not shared with any other subpopulation. Observed and expected heterozygosity are the proportion of loci that are heterozygous based on Hardy-Weinberg frequencies. π is a measure of nucleotide diversity. FIS indicates the inbreeding coefficient.

